# Awake ripples enhance emotional memory encoding in the human brain

**DOI:** 10.1101/2021.11.17.469047

**Authors:** Haoxin Zhang, Ivan Skelin, Shiting Ma, Michelle Paff, Michael A. Yassa, Robert T. Knight, Jack J. Lin

**Author notes:** Equal contribution (Haoxin Zhang, Ivan Skelin).

## Abstract

Intracranial recordings from the human amygdala and the hippocampus during an emotional memory encoding and discrimination task reveal increased awake sharp-wave/ripples (aSWR) after encoding of emotional compared to neutral stimuli. Further, post-encoding aSWR-locked memory reinstatement in the amygdala and the hippocampus was predictive of later memory discrimination. These findings provide electrophysiological evidence that post-encoding aSWRs enhance memory for emotional events.

## Main

Multiple mechanisms have been proposed to explain the prioritized encoding of emotional experiences^1–3^, including the neuromodulatory effects on plasticity and the interplay between the amygdala and the hippocampus^1,4,5^. Several studies have found memory reinstatement during the immediate post-encoding period to be predictive of later memory performance ^6,7^. Sharp-wave/ripples (SWRs) are transient hippocampal oscillations (80-150 Hz), associated with synchronous neural activation in the hippocampus and the amygdala^8,9^, and are implicated in the binding of anatomically distributed memory traces^10^. Behaviorally relevant reactivation of emotional memory occurs during aSWRs ^11^, and disruptions of post-experience aSWR interfere with memory utilization^12^. Based on these findings, we hypothesized that aSWRs occurring immediately after stimulus encoding (post-encoding) facilitate emotional memory discrimination through the coordinated hippocampal-amygdala memory reinstatement. Using intracranial electroencephalographic (iEEG) recordings in epilepsy patients during the performance of an emotional encoding and discrimination task, we first confirm reports of better discrimination memory for arousing stimuli^3^. Next, we demonstrate that the number of aSWR events immediately after encoding is associated with both stimulus-induced arousal and the accuracy of later discrimination. Finally, the coordinated memory reinstatement between the amygdala and the hippocampus during post-encoding aSWRs is predictive of later memory discrimination performance, with the amygdala reinstatement showing a directional influence on the hippocampal reinstatement. Together, these findings provide evidence for aSWRs-mediated memory reinstatement in the amygdala and hippocampus as a mechanism accounting for better remembering of emotional experiences.

We performed simultaneous iEEG recordings from the amygdala (*n*_*electrode*_ = 20) and the hippocampus (*n*_*electrode*_ = 17, Fig. 2a) in 7 human subjects, while performing an emotional memory encoding and discrimination task^13,14^ (Methods, Fig. 1a). During the encoding stage, subjects were presented with a stimulus (image; stimulus encoding) and asked to rate the stimulus valence as negative, neutral, or positive (post-encoding/response). During the retrieval stage, subjects were presented with one of the 3 types of stimuli - Repeats (identical), Lure (slightly different) or Novel (stimuli not seen during encoding) - and classified each stimulus as “New” or “Old.”

**Fig 1.**
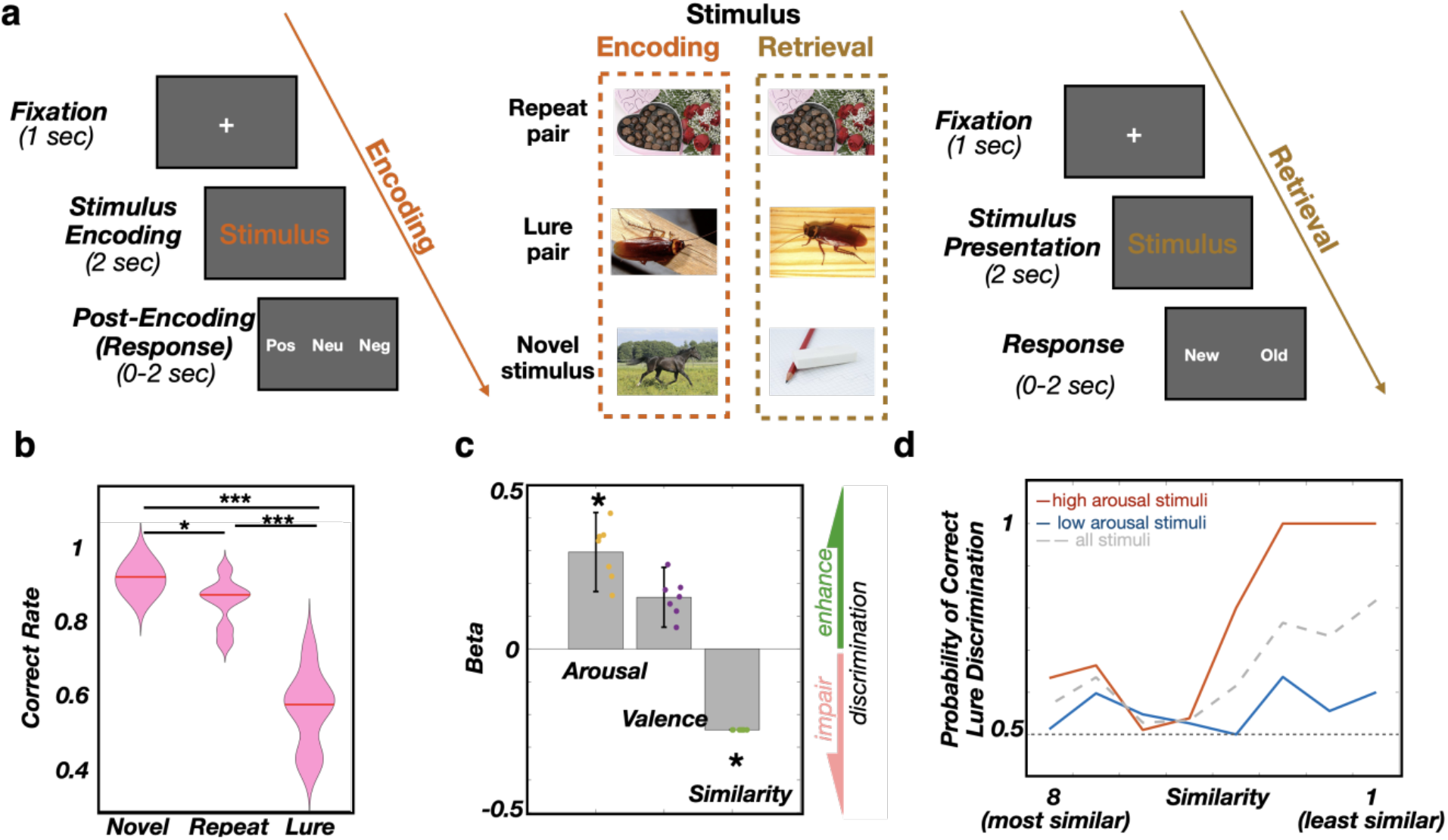
Memory discrimination is more accurate for emotional stimuli. **a**, Task structure: subjects are presented with an image (Stimulus encoding). Following presentation, they rate the valence of the image as negative, neutral, or positive (Post-Encoding/Response). Once all images are presented and rated, subjects are presented with 3 types of stimuli - Repeat (identical), Lure (slightly different) or Novel (stimuli not seen during encoding) - and classify each stimulus as “old” or “new.” **b**, Correct discrimination is highest for Novel stimuli (93.9 ± 1.4 %; median ± SEM), followed by Repeats (89.4 ± 2.4 %) and Lures (61.5 ± 3.7 %). Paired t-test: Novel vs. Repeat, *p = 0.016, t = 3.33, df = 6; Novel vs. Lure, ***p<0.001, t = 8.36, df = 6; Repeat vs. Lure, ***p < 0.001, t = 6.13, df = 6. **c**, Correct discrimination of Lure stimuli is positively associated with encoded stimulus-induced arousal (*p=0.047, β = 0.3 ± 0.12, t = 1.98, df = 452, logistic linear mixed-effect model) and valence (p = 0.137, β = 0.15 ± 0.09, t = 1.48, df = 452), while negatively associated with similarity (*p = 0.039, β = -0.24 ± 0.00, t = -2.06, df = 452). The β sign and magnitude indicate effect direction and strength, respectively. Dots correspond to individual subjects. **d**, Probability of Lure correct discrimination as a function of SI and stimulus-induced arousal. The solid line shows the actual proportion of ‘New’ responses (y-axis) as a function of Lure stimulus SI (x-axis) for low arousal (blue) or high arousal stimuli (red). The low/high arousal groups were created using the median split.

**Fig 2.**
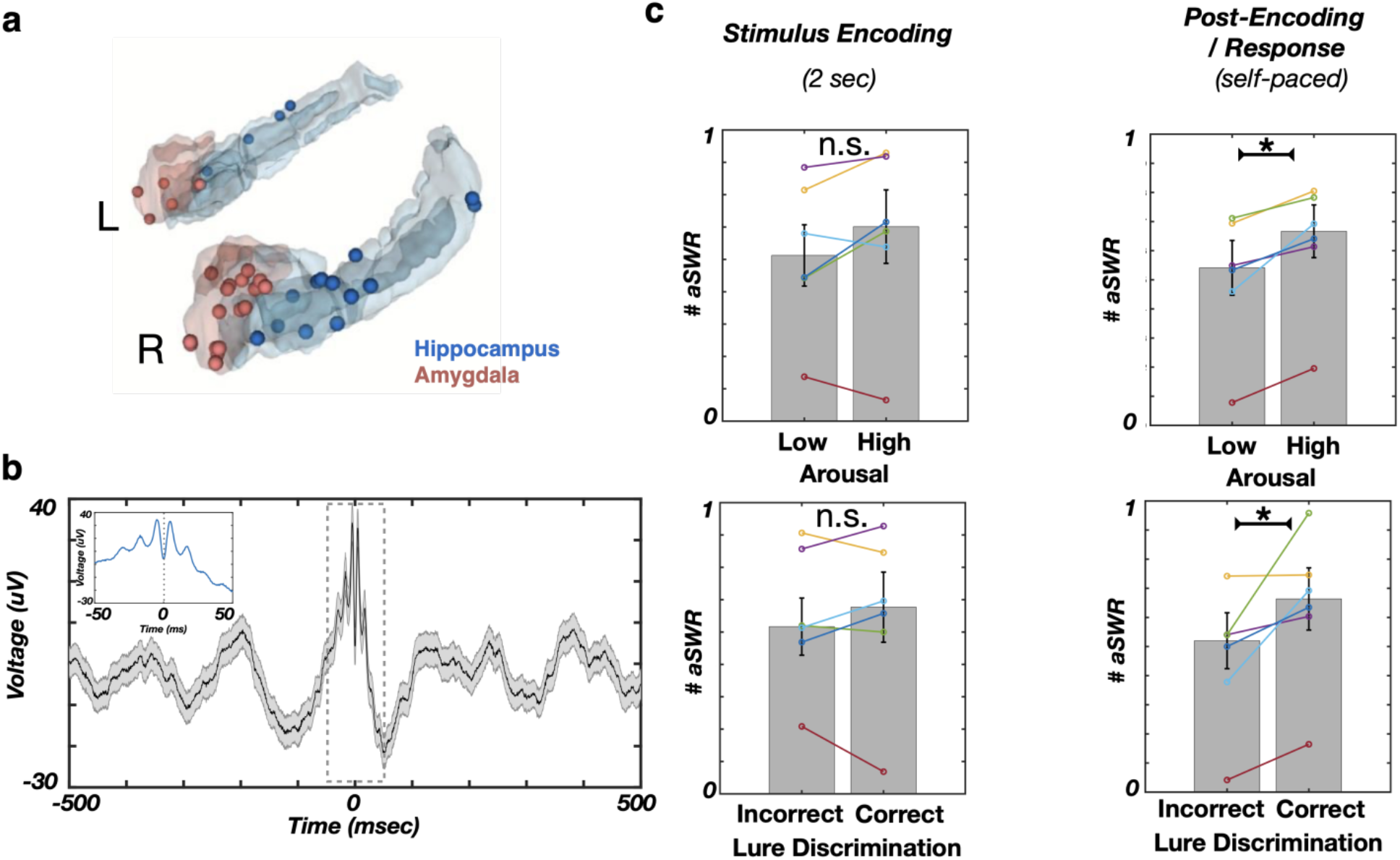
The post-encoding aSWR occurrence predicts the stimulus-induced arousal and memory discrimination. **a**, Reconstructed locations of hippocampal (blue) and amygdala electrodes (red). **b**, The aSWR grand average waveform (n = 4689 aSWRs in 6 hippocampal channels, 6 subjects). **c**, The aSWR occurrence is significantly higher following encoding of arousing (top right; *p = 0.03) and later correctly discriminated stimuli (bottom right, *p = 0.03). The aSWR occurrence was showing no conditional differences during stimulus encoding (left column, p’s > 0.05).

Memory discrimination is defined as the correct classification of: 1) Repeat stimuli as Old, 2) Novel stimuli as New, or 3) Lure stimuli as New. Subjects classified Repeat stimuli and Novel stimuli with high accuracy (Repeat: 89.4 ± 2.4%, Novel: 93.9 ± 1.4%; Fig. 1b). Memory discrimination accuracy was lower for Lure stimuli, relative to both Repeat or Novel stimuli (Lure: 61.5 ± 3.7 %; p_Novel vs Lure_ < 0.001, t = 8.36; p_Repeat vs Lure_ < 0.001, t = 6.13, paired t-test), reflecting similarity-induced memory interference. Indeed, there was a strong negative association between subjects’ stimulus discrimination ability and stimulus similarity rating (p = 0.039, t = -2.06, see Methods, Fig. 1c-d). Stimulus-induced arousal (irrespective of valence) was associated with better memory discrimination, confirming previous reports^1–3^ (p = 0.047, t = 1.98, Fig. 1c-d, Extended Data Fig. 1).

We defined the post-encoding period as the interval between stimulus offset and subjects’ stimulus valence rating response (Fig. 1a). We tested the association of post-encoding aSWR occurrence (i.e., the number of aSWRs) with the stimulus emotional content (stimulus-induced arousal and valence) and correct discrimination during retrieval. Higher post-encoding aSWR occurrence was associated with stimulus-induced arousal (p = 0.03, z = -2.2, Wilcoxon signed-rank test, Fig. 2c) and also predicted correct discrimination during retrieval (p = 0.03, z = -2.2, Wilcoxon signed-rank test, Fig. 2c), but was not associated with stimulus valence (p = 0.77, F(2, 15) = 0.25, one-way ANOVA; Extended Data Fig. 3). Taken together, these results provide the first report of post-encoding aSWRs as a potential electrophysiological mechanism for enhanced memory discrimination of arousing stimuli, previously characterized at behavioral level^2,3,15^. Furthermore, the positive associations between aSWRs and stimulus-induced arousal/later discrimination were present in all individual subjects (Fig. 2c). The post-encoding response time (RT) did not differ based on stimulus-induced arousal (p = 0.2, z = 0.7, RT_high-arousal_ = 0.8 ± 0.1 sec; RT_low-arousal_ = 0.6 ± 0.2 sec) or later discrimination (p = 0.25, z = 0.6, RT_correct_ = 0.7 ± 0.2 sec, RT_incorrect_ = 0.7 ± 0.3, Wilcoxon signed-rank test). Therefore, the associations between stimulus-induced arousal or correct discrimination and post-encoding aSWR occurrence were unrelated to post-encoding duration. Associations between aSWR and stimulus-induced arousal/later correct discrimination accuracy were selective for the post-encoding time window. These relationships were absent for the stimulus encoding or the retrieval task stage (p > 0.05, Wilcoxon signed-rank test; Fig. 2c, Extended Data Fig. 3, 4). The aSWRs probability was significantly higher during low theta power periods (Extended Data Fig. 5), consistent with observations that cholinergic tone promotes theta oscillations and suppresses SWRs^10,12^. In addition, aSWRs did not overlap with increased broadband gamma power, suggesting that aSWRs are distinct from non-specific broadband power fluctuations^16^ (Extended Data Fig. 5).

Recent studies suggest that post-encoding memory reinstatement supports successful subsequent memory retrieval ^6,7^. Meanwhile SWR is associated with reactivation of pre-established neuronal patterns^17^. We hypothesized that memory reinstatement during the post-encoding aSWR window could enhance later memory discrimination. Distinct neural populations have been proposed to represent individual stimuli, resulting in stimulus-specific high-frequency activity (HFA) patterns^18,19^. We, thus, quantified memory reinstatement as the Spearman correlation between HFA power spectral vectors (PSVs), for each combination of the encoding-response time bins from the same trial (Extended Data Fig. 6). Next, we computed the average reinstatement activity during ± 250 msec around post-encoding aSWR peaks. The reinstatement significance was determined relative to a null distribution, obtained by circular jittering of aSWR timestamps. The post-encoding aSWR-locked memory reinstatement was stronger for arousing and correctly discriminated stimuli (Extended Data Fig. 7). To assess specific contributions of the amygdala and the hippocampus to this phenomenon, we calculated post-encoding memory reinstatement for each region, relative to aSWR peak (Fig. 3a). The significant reinstatement period in the amygdala consisted of two intervals, the first starting slightly earlier and overlapping with the hippocampal reinstatement (−105 to -50 msec), and a second period following the hippocampal reinstatement (40 to 200 msec). The significant reinstatement period in the hippocampus lasted from -100 to 50 msec (Fig. 3b). These results demonstrated region-specific timing of the post-encoding aSWR-locked memory reinstatement in the amygdala and the hippocampus. Next, we tested for the temporal compression^17^ of post-encoding aSWR-locked reinstatement (no compression, 2x, 4x, and 6x compression) and showed the strongest aSWR-locked reinstatement with no compression (Extended Data Fig. 8). We then analyzed the association of the post-encoding memory reinstatement with the stimulus-induced arousal and later discrimination. Remarkably, we observed a region-specific double dissociation. Specifically, the amygdala, not the hippocampus, showed a positive association between aSWR-locked memory reinstatements and the stimulus-induced arousal (AMY: -80 to -10 msec, p = 0.035; HPC: p > 0.05, see Methods; Fig. 3c). In contrast, the hippocampus, but not the amygdala, revealed a positive association between aSWR-locked memory reinstatement and later correct discrimination (AMY: p > 0.05; HPC: -15 to 90 msec, p = 0.008, see Methods; Fig. 3c). To summarize, post-encoding aSWR-locked memory reinstatements in the amygdala and the hippocampus followed distinct temporal dynamics and were associated with reactivation of distinct aspects of encoded stimuli (i.e., the amygdala for stimulus-induced arousal and the hippocampus for later discrimination accuracy).

**Fig 3.**
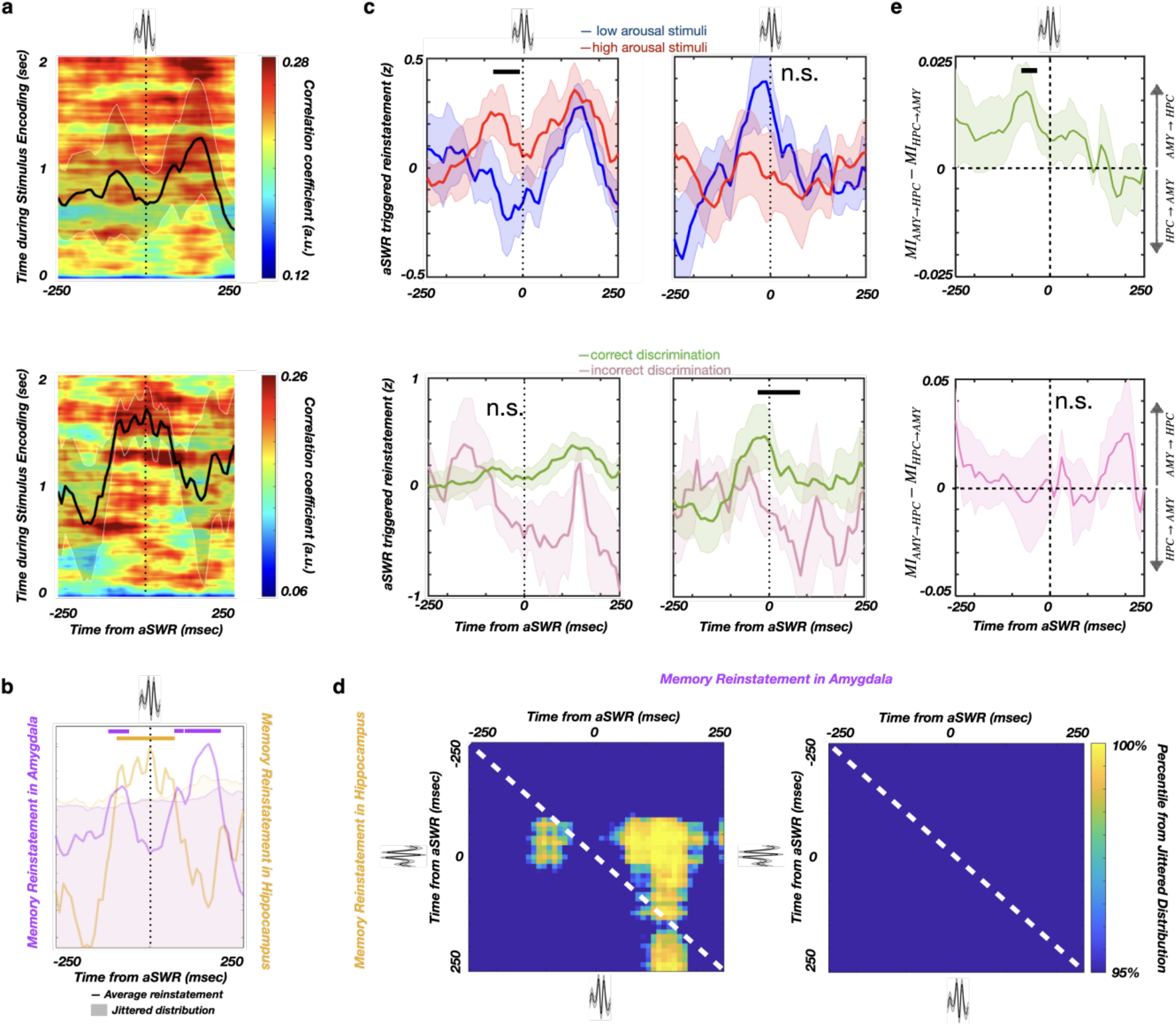
Memory reinstatement in the hippocampus and amygdala around aSWR. **a**, aSWR-locked reinstatement in the amygdala (top) and hippocampus (bottom) during the post-encoding period (line and shaded areas represent the mean ± SEM). **b**, Reinstatement is greatest around the time of aSWRs as shown by comparison with the null-distribution (within ± 250 msec). Shaded areas denote the null-distribution 95% confidence interval. Reinstatement in the hippocampus overlaps with aSWR peak (orange), while reinstatement in the amygdala peaks prior to and after the aSWR (magenta). **c**, aSWR-locked reinstatement in the amygdala is stronger for arousing stimuli (top left, p = 0.035, see Methods) but is not associated with subsequent discrimination (bottom left, p = 0.066). Reinstatement in the hippocampus is robust for correctly discriminated stimuli (bottom right, p = 0.008, see Methods) but does not depend on stimulus-induced arousal (top right, p > 0.1). **d**, The aSWR-locked joint reinstatement in the hippocampus and amygdala for the correct (left) and incorrect (right) discrimination trials. Reinstatement in the amygdala starts 100 msec prior to the aSWR peak, followed by reinstatement in the hippocampus (−50 to 200 msec). There is no significant joint reinstatement during incorrect discrimination trials, suggesting that the cross-structure joint reinstatement may be required for correct discrimination. **e**, Mutual information (MI) difference for the amygdala (AMY) and hippocampal (HPC) memory reinstatement time-courses, during the post-encoding aSWR windows (correct discrimination - top, incorrect discrimination - bottom). Positive values denote stronger AMY→HPC directionality. A temporal cluster of significant MI difference (AMY→HPC) is present before aSWR peak time(-70 to -30 msec) after encoding of correctly discriminated stimuli (top; p = 0.038, see Methods), indicating that hippocampal reinstatement is better predictable by amygdala reinstatement than vice versa. This effect is present only during the post-encoding period for correctly discriminated stimuli (top), but not for the incorrectly discriminated stimuli

In rodents, the coordinated memory reactivation in the amygdala and hippocampus during sleep SWRs is proposed to bind neuronal ensembles encoding emotional and spatial information, respectively^20^. We reasoned that a similar interaction between the amygdala and the hippocampus exists in which cross-regional post-encoding aSWR-locked memory reinstatement facilitates later discrimination. We hypothesized that the reinstatement in both structures co-occurs during the same aSWR events and follows a consistent temporal dynamic. To test this, we separately computed aSWR-locked joint memory reinstatement for the correctly and incorrectly discriminated stimuli (Methods). A significant joint aSWR-locked memory reinstatement in the amygdala and hippocampus was present during the post-encoding period only for correctly discriminated stimuli (Fig. 3d; Extended Data Fig. 9). Specifically, the amygdala reinstatement preceded the hippocampal reinstatement by ∼100 msec. Further, mutual information analysis showed a significant unidirectional influence from the amygdala to the hippocampus before aSWR peak (−70 to -30 msec, p = 0.038; see Methods; Fig. 3e). To conclude, aSWR-mediated coordination of memory reinstatement in the amygdala and the hippocampus promotes later successful discrimination.

Rodent studies have implicated the SWRs in the retrieval and consolidation of emotional memory. However, it is unclear whether it supports the memory benefits of emotional experience^21^. Our study reveals an association of higher aSWR occurrence with stimulus-induced arousal and subsequent correct stimulus discrimination, providing direct evidence for aSWR-mediated strengthening of emotional memory. Interestingly, the higher aSWRs occurrence has been shown in rodents, after exposure to a novel or reward-associated context^22^. Together, this suggests that aSWRs may play a general role in the selective enhancement of salient experiences^23^.

Notably, such association is specific to the post-encoding period that starts immediately after memory encoding, when memory retrieval is essential to rate the emotional content of the stimuli. This finding supports theoretical assumptions that SWRs mediate both the retrieval of stored representation utilized in decision-making, and the strengthening of the same representation, contributing to memory consolidation^22^.

Next, we aimed to discern the link between the aSWR-associated interaction between the amygdala and hippocampus during post-encoding and subsequent memory effect. We found the aSWRs were accompanied by memory reinstatement during the post-encoding period. Specifically, the reinstatement in the amygdala appears shortly before the aSWR peak and shows association with arousing stimuli, while the hippocampal reinstatement appears around the aSWR peak and shows associations with correct subsequent memory discrimination. Moreover, the co-occurrence of the amygdala and the hippocampal reinstatement during the same post-encoding aSWR events - with the amygdala reinstatement leading hippocampal by ∼100 msec - is predictive of subsequent correct memory discrimination. This finding suggests that the coordinated reinstatement in the amygdala and hippocampus during aSWR is responsible for combining emotional and contextual aspects of the memory^20,21^.

Both the joint-reinstatement and mutual information analyses further confirm the predictive validity of directional influence from the amygdala to the hippocampus before aSWRs on correct discrimination, establishing a link between the amygdala reinstatement and memory discrimination as a physiological mechanism of emotional memory enhancement. Together, our data support a model wherein the memory reinstatement in the amygdala, triggered by emotional stimuli, elicits amygdala-hippocampal aSWR-associated memory reinstatement, enabling the coordinated joint-reinstatement, which facilitates subsequent memory performance.

## Acknowledgement

The authors thank all the participants for taking part in the study, as well as the nurses, technicians, and physicians at the UCI Epilepsy Unit. This work was supported by NIH Grant 1U19NS107609-01 to R.T.K. (subcontract to J.J.L).

## Extended data

**Extended Data Fig 1.**
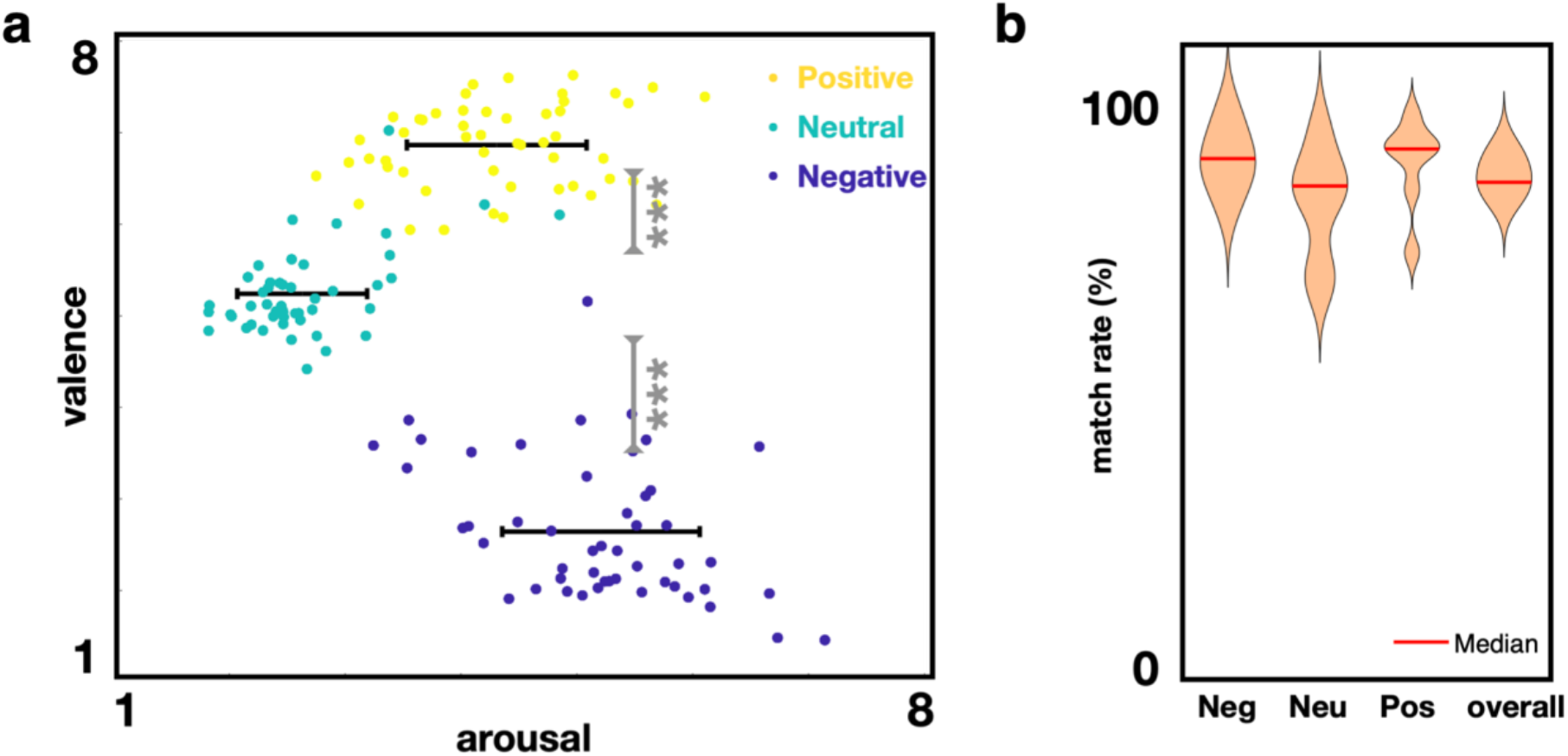
**a**, Positive and negative valenced stimuli are associated with higher stimulus-induced arousal, relative to neutral valence stimuli (***p<0.001, Wilcoxon rank-sum test). **b**, Stimuli valence ratings of study subjects are highly similar to the healthy population (match rate = 85.3 ± 1.3%).

**Extended Data Fig 2.**
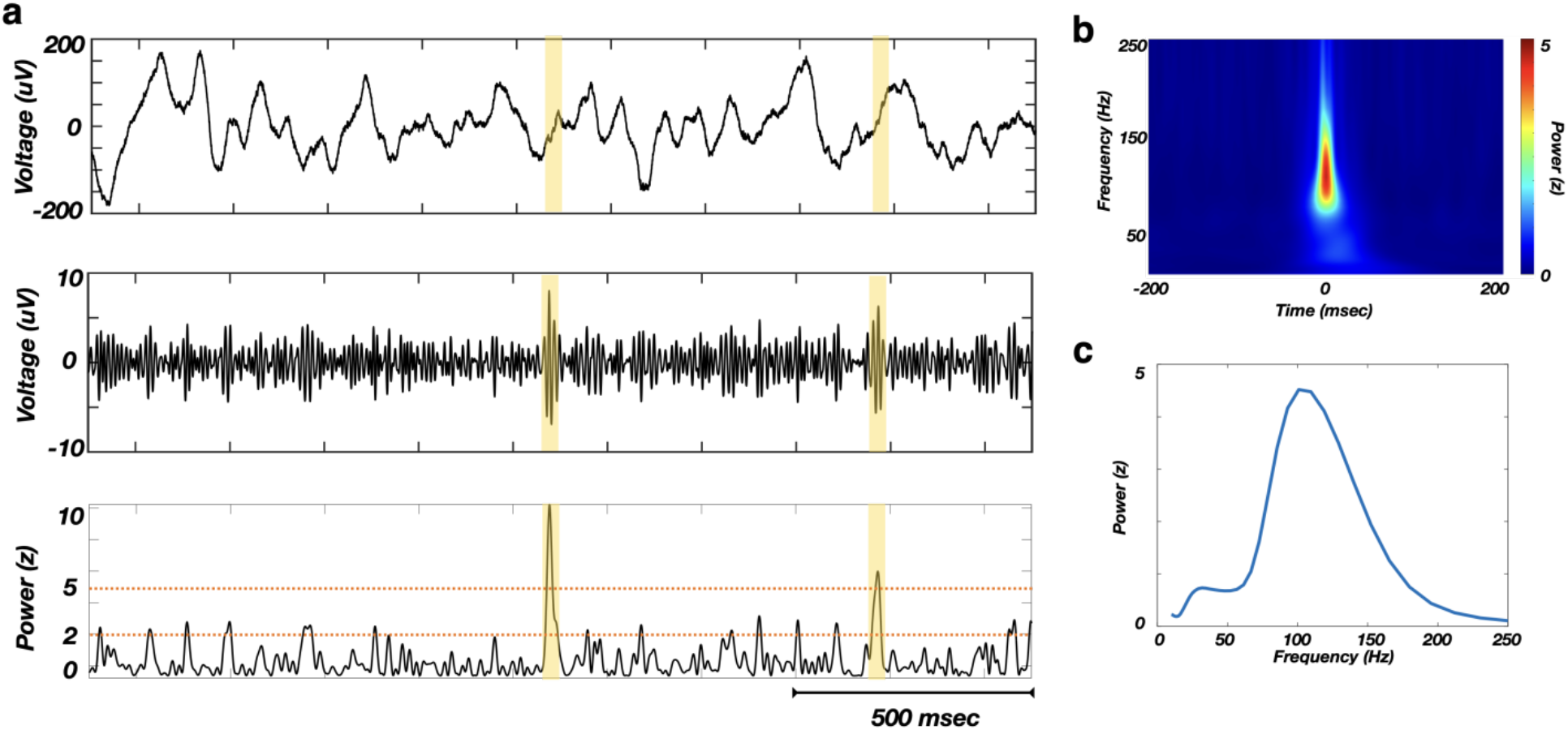
Awake SWR detection. **a**, Examples of several detected aSWRs (yellow highlights), showing the raw trace (top), filtered trace (80 - 150 Hz range, middle) and z-scored envelope of filtered trace (bottom). Detection is based on double-threshold (orange dashed lines) crossing of z-scored power (80-150 Hz) for the period of 20-100 msec. **b**, Z-scored power spectral density of average detected aSWR. **c**, Z-scored power during aSWR windows shows a bump in the 80-150 Hz range. This suggests that the aSWRs are not detected during signal artifact periods, which would reflect as a broadband power increase. In addition, detected aSWRs are not detected during non-specific increase in broadband gamma power or pathological high-frequency oscillations (> 200 Hz).

**Extended Data Fig 3.**
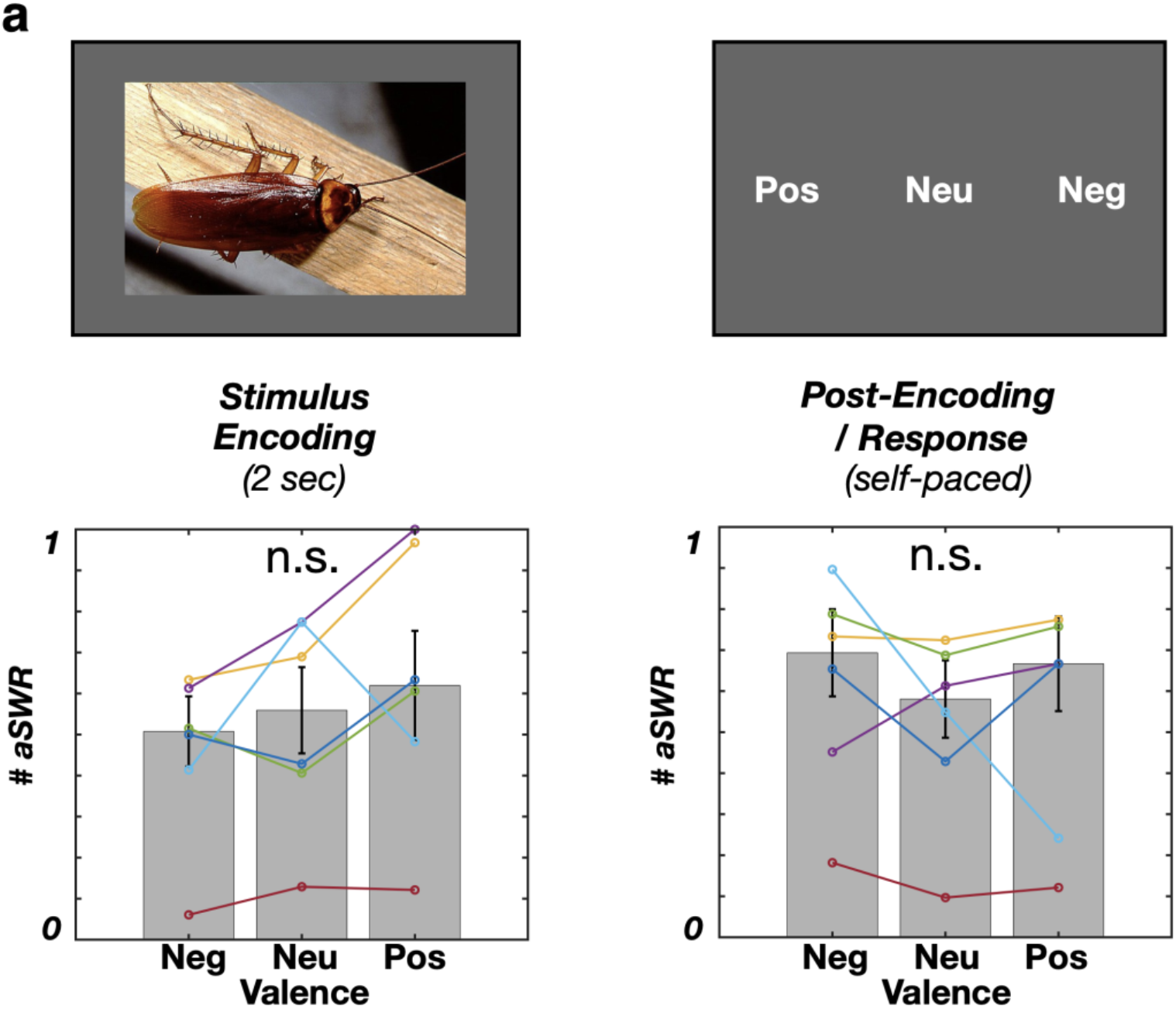
Stimulus valence is not significantly associated with aSWR occurrence during encoding stage. **a**, Stimulus encoding phase: F(2, 15) = 0.67, p = 0.53; Post-encoding: F(2, 15) = 0.25, p = 0.77, One-way ANOVA). The data from individual subjects are color-coded.

**Extended Data Fig 4.**
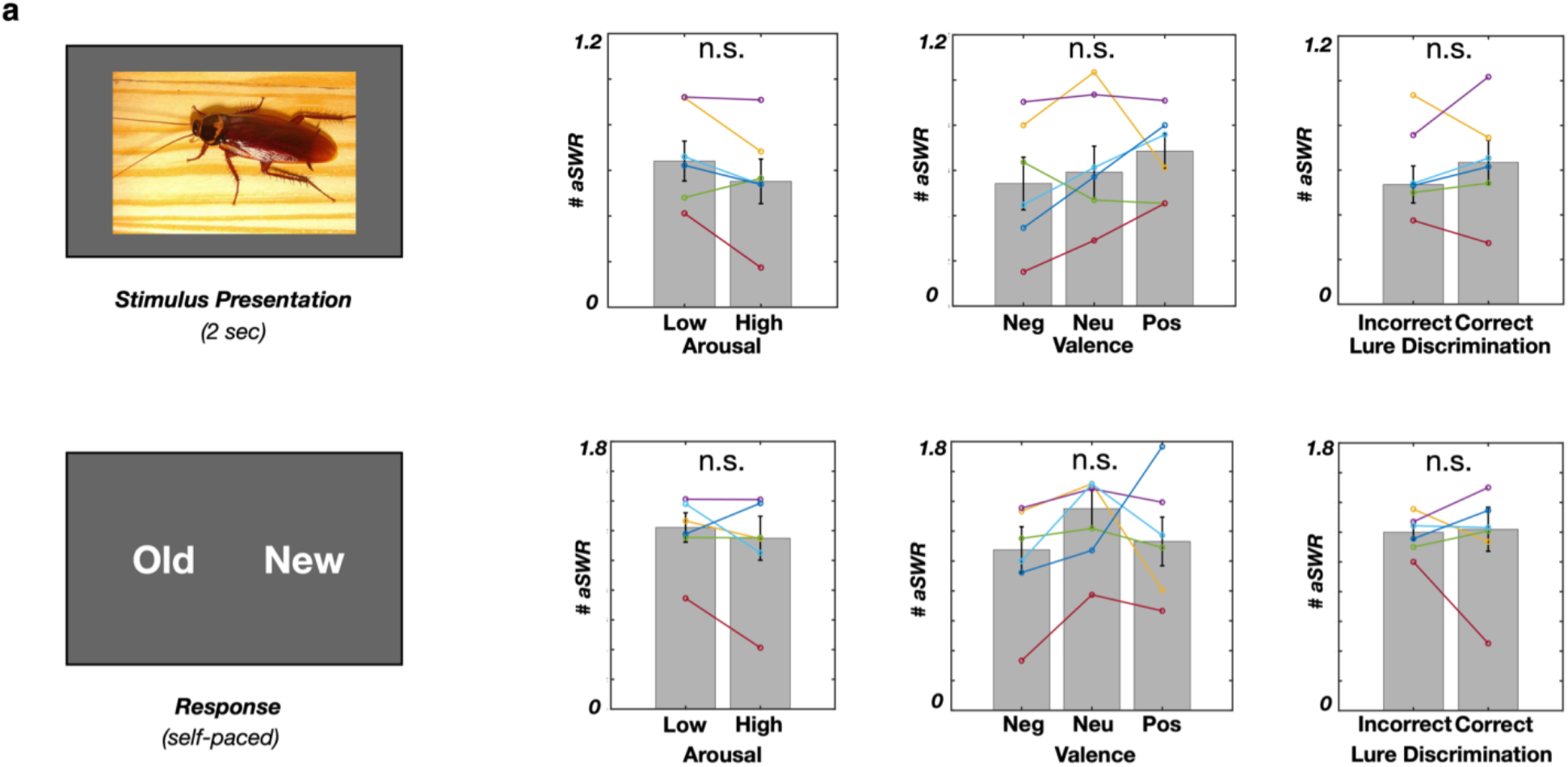
The aSWR occurence during retrieval task stage is not associated with stimulus-induced arousal, valence or correct discrimination. Arousal: Stimulus presentation (top row), p = 0.11, z = 1.57; Response (bottom row), p = 0.17, z = 1.36, Wilcoxon signed-rank test. **Valence**: Stimulus presentation, p = 0.69, F(2, 15) = 0.69; Response, p = 0.51, F(2, 15) = 0.71, One-way ANOVA). **Correct discrimination**: Stimulus presentation: p = 0.6, z = -0.52; Response: p = 0.92, z = 0.11, Wilcoxon signed-rank test).

**Extended Data Fig 5.**
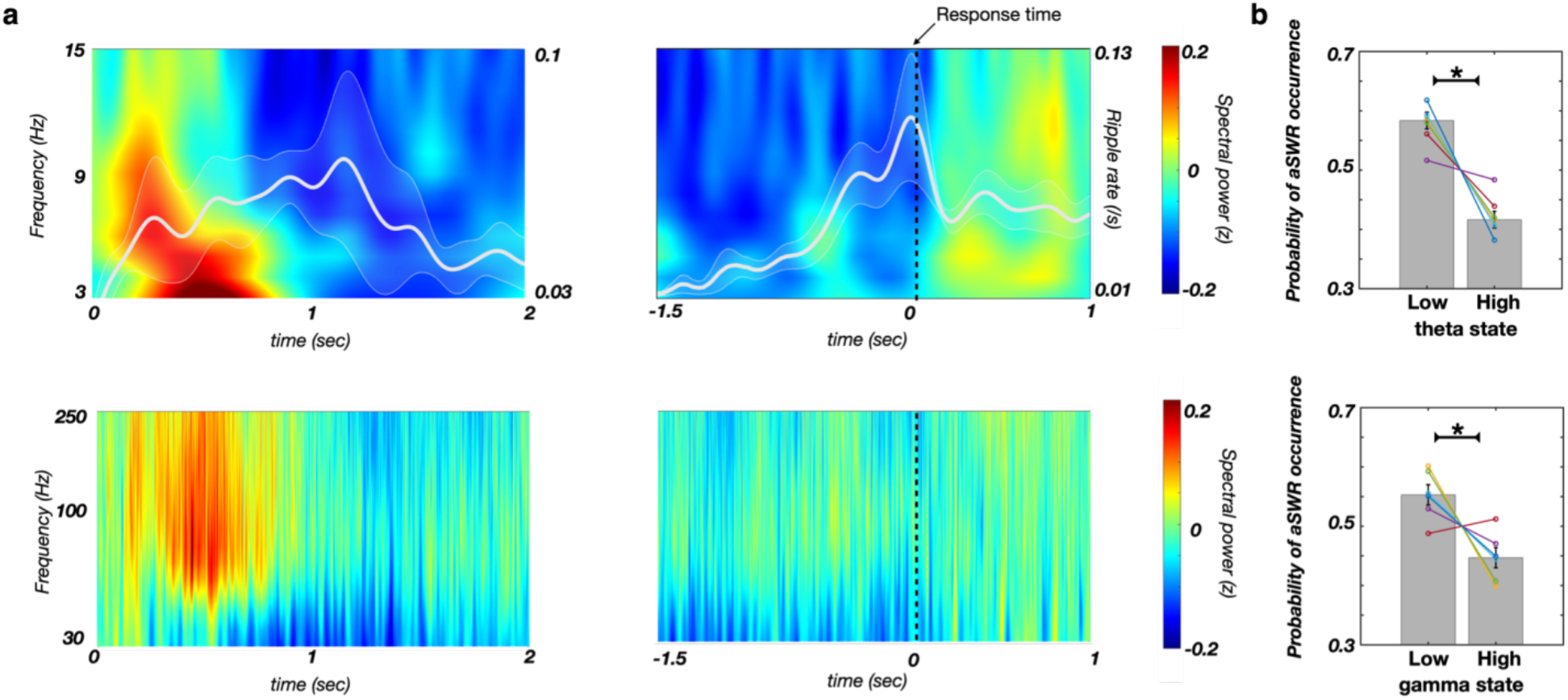
aSWRs occur predominately outside of high theta or broadband gamma periods. **a**, Low frequency (top, color) and high frequency spectrogram (bottom, color), and aSWR rate (white line) during the stimulus encoding (left) and post-encoding (right, response-locked) periods. **b**, The probability of aSWR occurrence is lower during the high theta state (top, p = 0.017, z = 2.1, one-tailed Wilcoxon signed-rank test), or during high gamma state (bottom, p = 0.028, z = 1.9, one-tailed Wilcoxon signed-rank test). Theta/gamma state classification was based on the power median split (for details, see ‘Dual state analysis’).

**Extended Data Fig 6.**
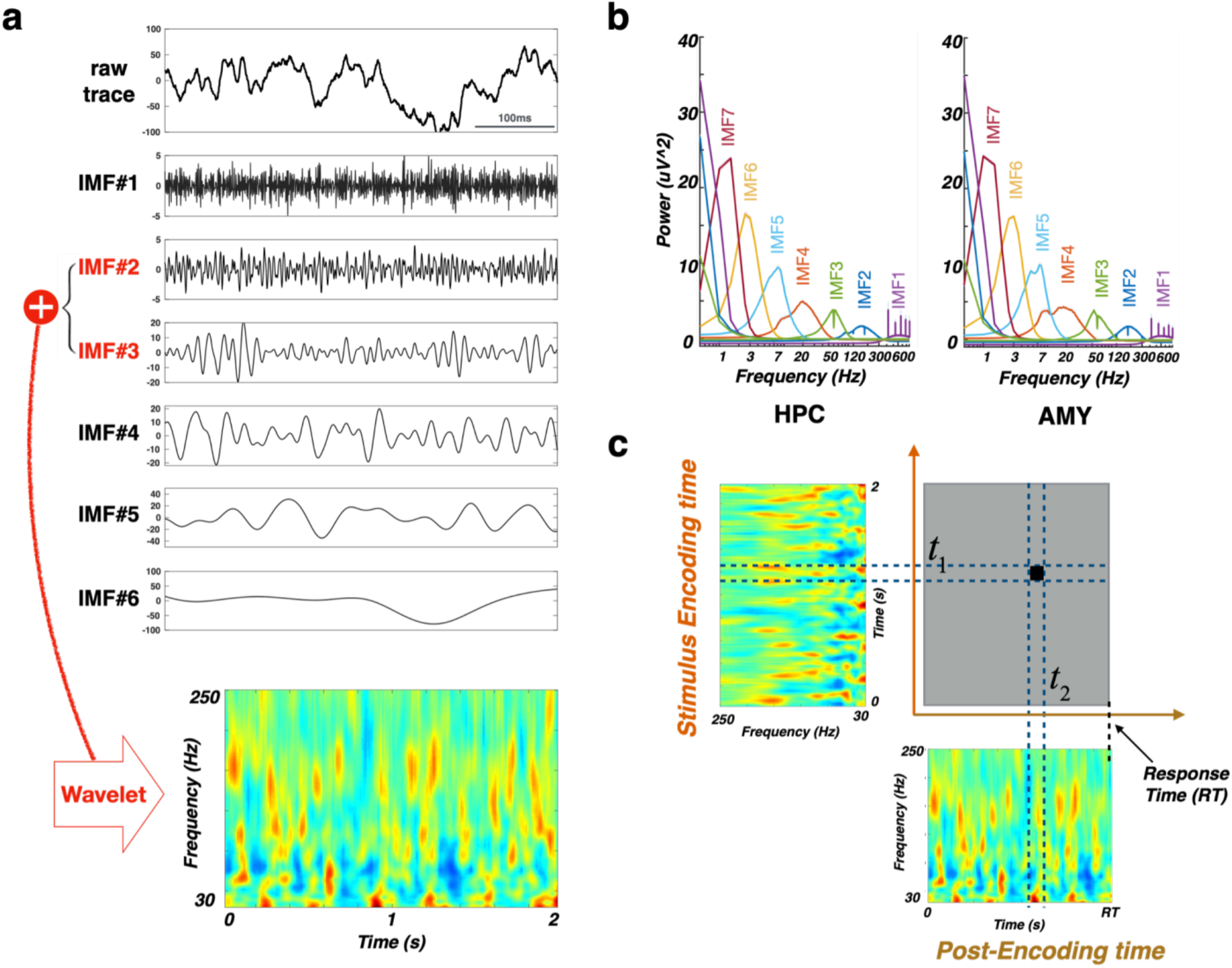
Overview of Ensemble Empirical Mode Decomposition (EEMD) and representational similarity analysis (RSA) methods. **a**, An example hippocampal raw iEEG trace (top) was decomposed into multiple intrinsic mode functions (IMFs; lower 6 panels). IMFs within the HFA range (IMF_2_ and IMF_3_) were used for HFA reconstruction. The HFA time-frequency matrix (bottom) was estimated using wavelet transformation (for details, see Time-frequency representation of the HFA). **b**, Power spectral density (mean ± SEM) of the IMFs decomposed from the hippocampal (left) and amygdala (right) electrodes. IMF spectral features were consistent across subjects and structures, with mean center frequencies in delta (IMF_7_), theta (IMF_6_, IMF_5_), alpha/beta (IMF_4_), gamma (IMF_3_), high-gamma bands (IMF_2_), and the noise term (IMF_1_). The HFA time series were estimated by summing the IMFs with center frequencies > 30 Hz (IMF_2_ and IMF_3_). **c**, The similarity matrix (top right) was constructed by computing the power spectrum vector (PSV) Spearman’s correlations for each combination of stimulus encoding (top left) and post-encoding (bottom right) time bins.

**Extended Data Fig 7.**
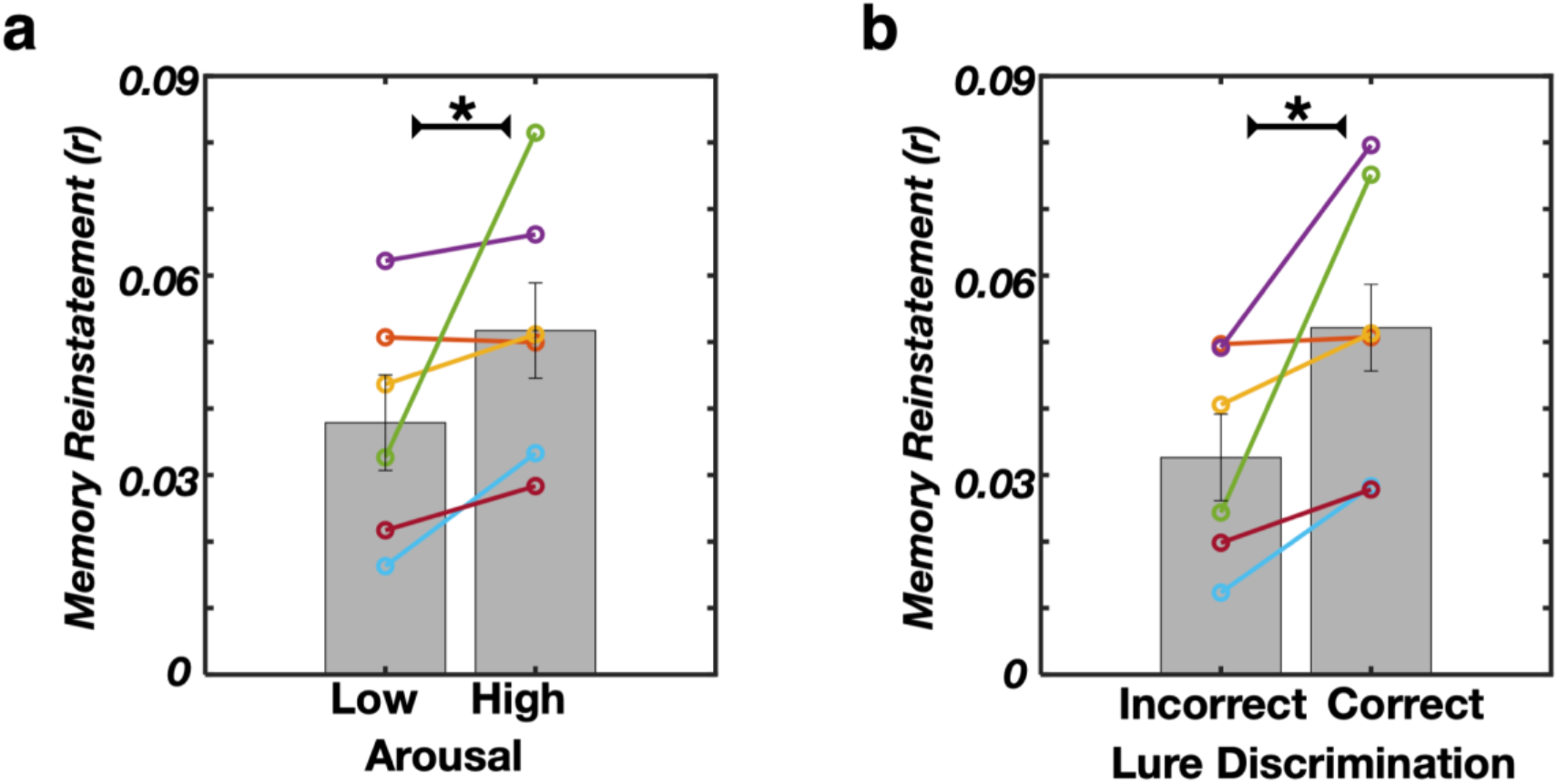
Post-encoding aSWR-locked reinstatement (amygdala and hippocampus combined) is increased for high stimulus-induced arousal and correctly discriminated stimuli. **a**, Arousal: *p = 0.046, z = -1.991, Wilcoxon signed-rank test. **b**, Correct discrimination: *p = 0.028, z = -2.201, Wilcoxon signed-rank test). Data from individual subjects is color-coded.

**Extended Data Fig 8.**
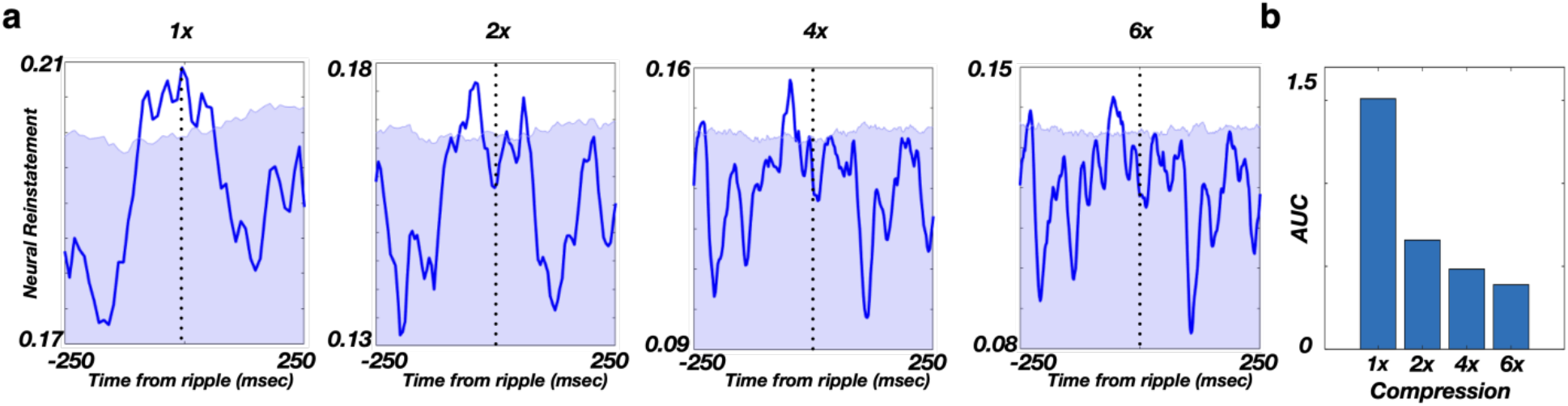
Post-encoding aSWR-locked memory reinstatement in the hippocampus is strongest without time compression. **a**, aSWR-locked hippocampal reinstatement during the post-encoding response period, across different temporal compression factors. Memory reinstatement strength area under curve (AUC) is defined as enclosed by the reinstatement trace (blue line) and 95% percentile of empirical null-distribution (blue shading upper limit). AUC reflects the memory reinstatement strength at different compression factors. **b**, Memory reinstatement strength is highest with no compression.

**Extended Data Fig 9.**
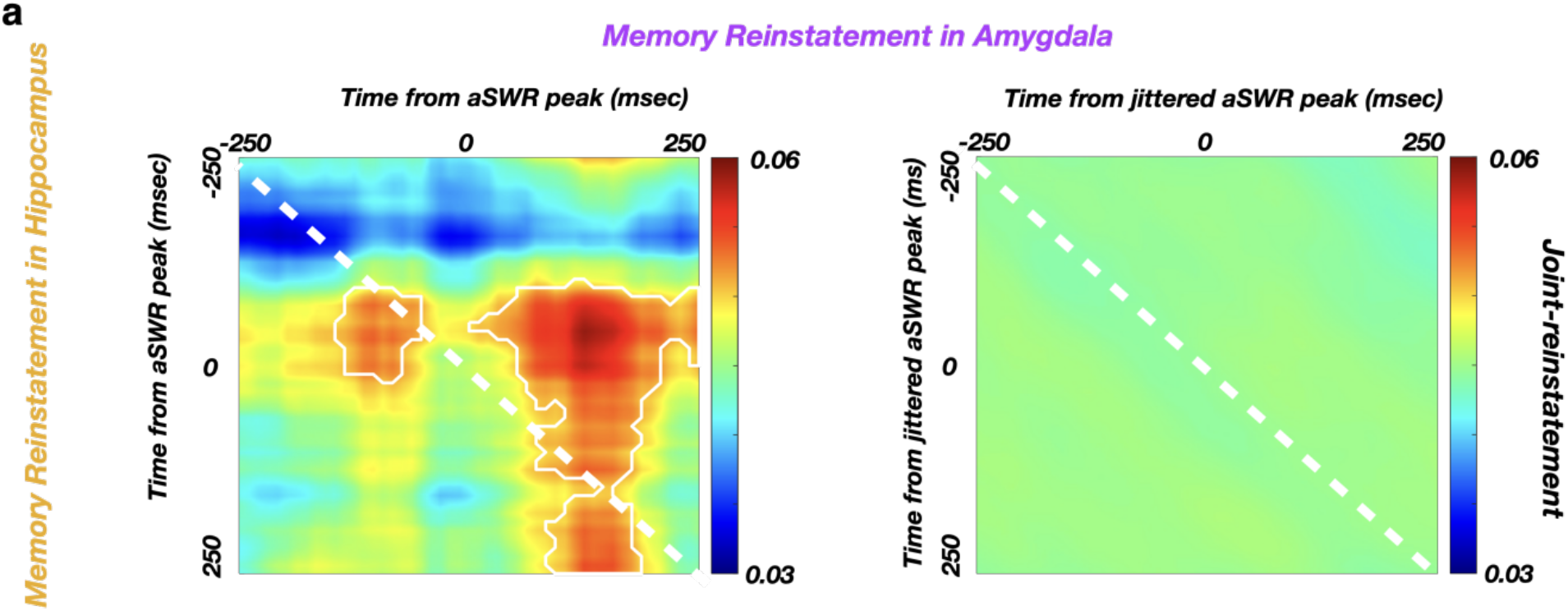
Joint cross-structure memory reinstatement occurs selectively during aSWR time windows. **a**, Average joint cross-structure reinstatement (hippocampus and amygdala) relative to aSWR peak times (left) and relative to jittered aSWR peak times (right). The white line encircles the periods of significant joint cross-structure memory reinstatement (Fig. 3d). The color scale represents the Spearman correlation between the encoding stimulus presentation and post-encoding aSWR windows. The absence of significant joint cross-structure memory reinstatement following the jittering of aSWR peak times (right) reveals the specificity of cross-structure reinstatement to aSWR windows.

**Extended Data Table 1.**
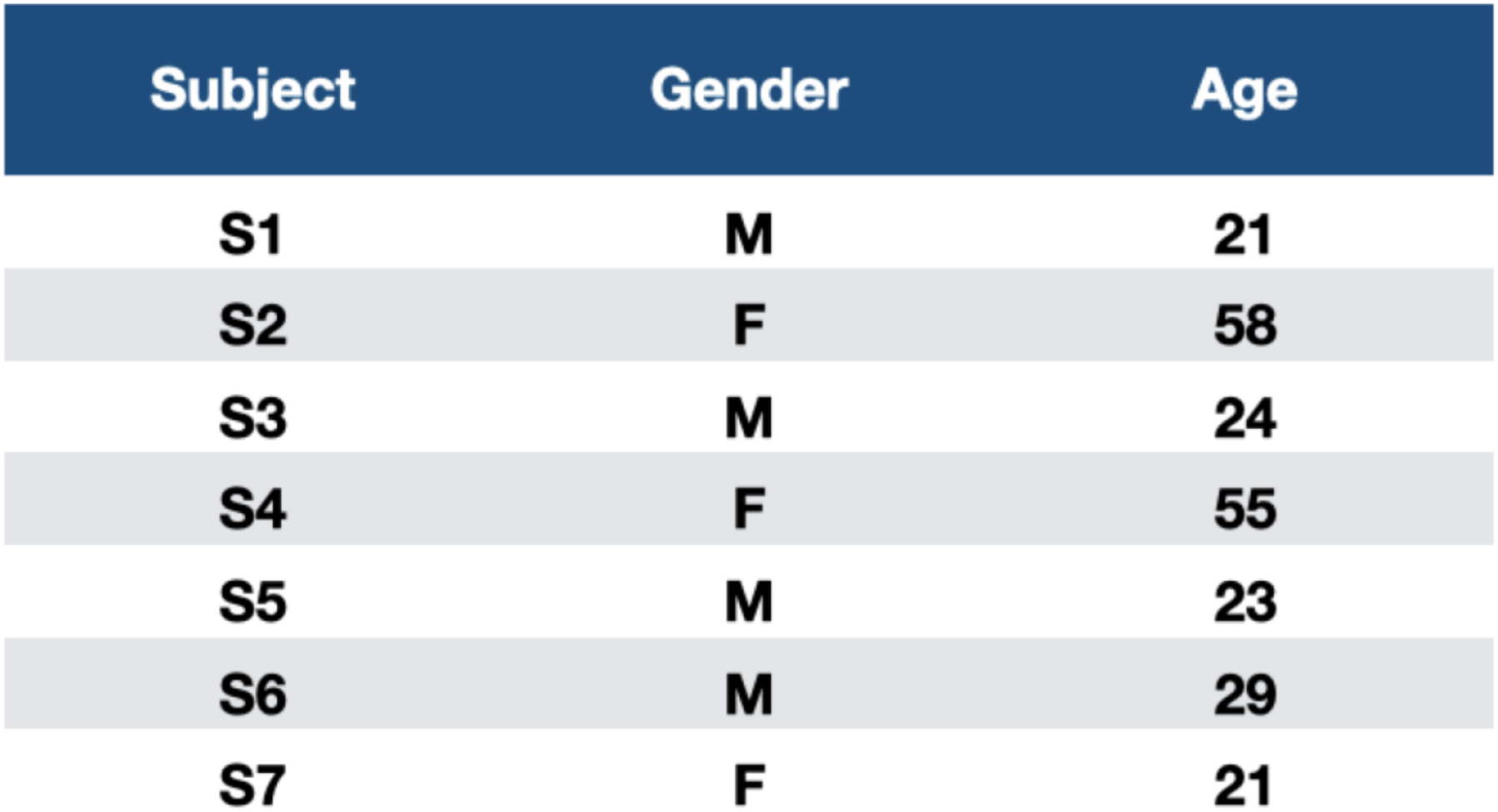
Demographic information for the study subjects.

| Subject | Gender | Age |
| --- | --- | --- |
| S1 | M | 21 |
| S2 | F | 58 |
| S3 | M | 24 |
| S4 | F | 55 |
| S5 | M | 23 |
| S6 | M | 29 |
| S7 | F | 21 |

**Extended Data Table 2.**
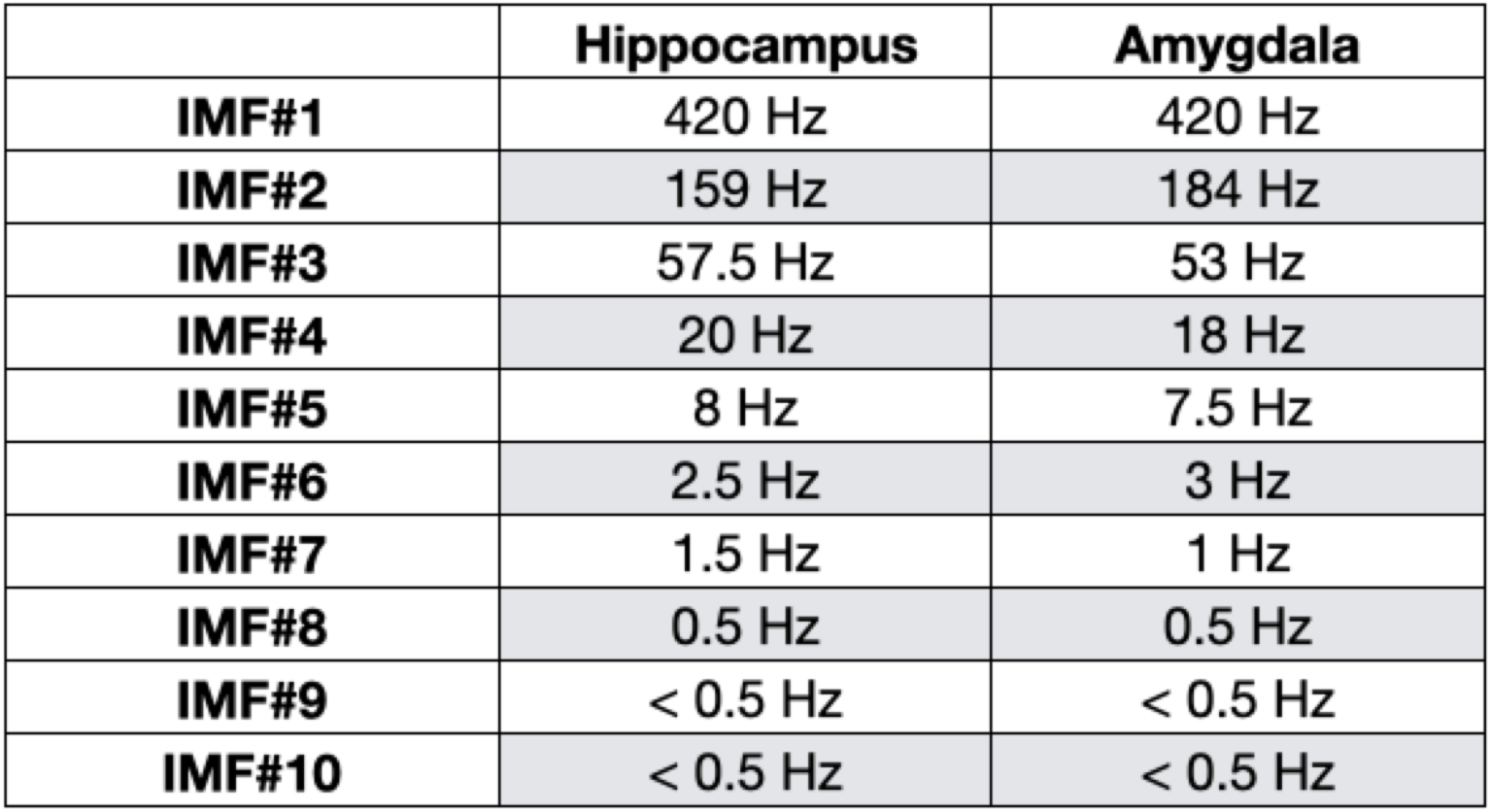
Center frequencies of the IMFs in the hippocampus and amygdala.

|  | **Hippocampus** | **Amygdala** |
| --- | --- | --- |
| **IMF#1** | 420 Hz | 420 Hz |
| **IMF#2** | 159 Hz | 184 Hz |
| **IMF#3** | 57.5 Hz | 53 Hz |
| **IMF#4** | 20 Hz | 18 Hz |
| **IMF#5** | 8 Hz | 7.5 Hz |
| **IMF#6** | 2.5 Hz | 3 Hz |
| **IMF#7** | 1.5 Hz | 1 Hz |
| **IMF#8** | 0.5 Hz | 0.5 Hz |
| **IMF#9** | < 0.5 Hz | < 0.5 Hz |
| **IMF#10** | < 0.5 Hz | < 0.5 Hz |

## Methods

### Subjects

Intracranial electroencephalography (iEEG) recordings were obtained from 7 subjects (3 females; mean age ± SD = 33 ± 16), undergoing presurgical monitoring of epileptic foci at the University of California Irvine Medical Center (UCIMC) Epilepsy Monitoring Unit. The individual subject demographic information is shown in Table 1. Only the subjects with the correct discrimination rate of Novel trials >= 85% (see Emotional memory encoding and discrimination task) were included in the analysis. Electrode placements were determined entirely based on clinical considerations. All the research procedures were approved by the UCI Institutional Review Board and data was collected following informed consent.

### Statistics

All the statistical tests were performed with the individual subject as the unit of analysis. Unless stated otherwise, all the parametric statistical tests (e.g., Wilcoxon signed-rank test, t-test) were two-tailed. The effects of valence, stimulus-induced arousal and similarity on stimulus discrimination (Fig. 1c) were assessed using the logistic linear mixed-effect model (for details, see Behavioral Analysis). Conditional comparisons of aSWR occurrence (correct/incorrect discrimination or high/low arousal; Fig. 2c) were done using the Wilcoxon signed rank test (p < 0.05). Statistical significance of aSWR-locked memory reinstatement strength (Fig. 3b) was assessed by comparing the real test statistics with empirical null distribution, obtained using Monte Carlo method (for details, see Representational Similarity Analysis). We implemented the cluster-based nonparametric permutation test^1^ to assess the conditional differences (correct/incorrect discrimination or high/low arousal) of memory reinstatement strength (Fig. 3c), mutual information (Fig. 3e), by randomly shuffling the conditional trial labels 1000 times (for details, see Representational Similarity Analysis).Similarly, the significant temporal windows for the cross structure aSWR-locked joint memory reinstatement (Fig. 3d) were assessed by comparing to empirical null distribution (for details, see Joint-reinstatement Analysis).

### Emotional memory encoding and discrimination task

The emotional memory encoding and discrimination (EMOP) task consists of encoding and discrimination blocks. During the encoding block (148 trials), each trial consists of a cross fixation (1000 msec), followed by stimulus encoding (2000 msec) and self-paced post-encoding response period (up to 2000 msec). During the post-encoding response period, subjects are asked to classify the stimulus emotional valence as either negative, neutral or positive, using the corresponding laptop key. During the retrieval block (290 trials), trial time structure is identical to encoding phase. Following the cross fixation (1000 msec), the subjects are presented for 2000 msec with a stimulus identical (Repeat, 54 trials), slightly different (Lure, 97 trials) or unrelated (Novel, 139 trials) to previously encoded stimuli. Next, during the self-paced memory discrimination epoch (up to 2000 msec), subjects are asked to discriminate if the presented stimulus was seen during encoding (Old) or not (New). Correct discrimination is defined as classifying the Repeat stimuli as Old and Lure or Novel stimuli as New. The stimuli were selected from the continuous distributions across the valence and stimulus-induced arousal axes (Extended Data Fig. 1). The same set of stimuli was used across subjects. In addition, the valence, arousal and similarity of each stimulus were rated by separate cohorts of healthy subjects. Specifically, a first cohort (*N* = 50, 32 females; age mean ± SD = 22 ± 5) rated the stimulus emotional valence on a continuous scale (range 1-9, with 1 denoting the most negative, 9 the most positive, and 5 neutral valence). Stimuli were assigned in Negative (valence <= 3.5), Neutral (3.5 > valence < 6) or Positive (valence >= 6) groups. Another cohort of healthy subjects (*N* = 16, 4 females; age mean ± SD = 23 ± 5) rated the stimulus-induced emotional arousal on a scale 1 - 9 (1 being the least and 9 being the most arousing). Finally, a third cohort (*N* = 17, 11 females; age mean ± SD = 20 ± 1) examined relative similarity on the scale 1-8 ^2^. The high correspondence of stimulus valence ratings obtained from study subjects and healthy population (match rate = 85.3 ± 1.3%) suggests the intact emotional processing in study subjects (Extended Data Fig. 1).

### Behavioral Analyses

To assess the effects of valence, stimulus-induced arousal and similarity on Lure stimulus discrimination, we implemented the logistic linear mixed-effect model

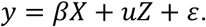

In this model, *y* indicates the responses across the individual Lure discrimination trials (0-Old; 1-New), *X* = [*x*_1_ *x*_2_ *x*_3_]^*T*^ denotes three fixed effect regressors (encoded stimulus valence and arousal as well as similarity between the encoded and Lure stimulus), *Z* = [*z*_1_]^*T*^ denotes random effect regressor (subject identity), *β*and *u* denote the fixed and random-effect regression coefficients, and *ε* denotes the error term. The model includes random intercept to incorporate individual subject differences. We normalized the valence, stimulus-induced arousal and similarity values relative to the scale of 0 to 1. The statistics reported in Fig. 1c corresponds to the fixed-effect coefficients *β*.

### Data collection

The behavioral experiment was administered using the PsychoPy2 software^3^ (Version 1.82.01). The laptop was placed at a comfortable distance in front of the subject. The iEEG signal was recorded using a Nihon Kohen system (256 channel amplifier, model JE120A), with an analog high-pass filter (0.01 Hz cutoff frequency) and sampling frequency 5000 Hz.

### Electrode localization

We localized each electrode using pre-implantation structural T1-weighted MRI scans (pre-MRI) and post-implantation MRI scans (post-MRI) or CT scans (post-CT). Specifically, we co-registered pre-MRI and post-MRI (or post-CT) scans by means of a rigid body transformation parametrized with three translation in x,y,z directions as well as three rotations using Advanced Normalization Tools (ANTs https://stnava.github.io/ANTs/). We implemented a high-resolution anatomical template with the label of medial temporal lobe subfields^2^ to guide the localization for individual electrodes. We resampled the template with 1mm isotropic, and aligned it to pre-MRI by ANTs Symmetric Normalization^4^ to produce a subject-specific template. The electrode localization was identified by comparing the subject-specific template subfield area with electrode artifacts.(Fig. 2a) The localization results were further reviewed by the neurologist (J.J.L.).

### Preprocessing

The signal preprocessing was done using the custom-written MATLAB code (Version 9.7) and Fieldtrip Toolbox^5^. The 60 Hz line noise and its harmonics were removed using a finite impulse response (FIR) notch filter (ft_preprocessing.m function in FieldTrip). The EEG signal was down-sampled to 2000 Hz, demeaned and high-passed filtered (cutoff frequency 0.3 Hz). The power spectrum density (PSD) was computed using the multitaper method with the Hanning window (ft_freqanalysis.m function in FieldTrip). All the channels were re-referenced to the nearest white matter channel from the same depth electrode, based on the electrode localization results. The interictal epilectic discharges were manually marked by an epileptologist (J.J.L.), using the ft_databrowser.m function in FieldTrip. The channels with severe contamination and trials containing epileptiform discharges were excluded from further analyses.

### Awake sharp-wave/ripple detection

Following the removal of channels with excessive epileptic activity and individual trials containing visually identified interictal epilectic discharges, awake sharp-wave/ripples (aSWRs) were detected on the remaining hippocampal channels, using the Freely Moving Animal Toolbox (FMA; http://fmatoolbox.sourceforge.net/). First, the iEEG traces from the trials used in the analysis were concatenated. Next, concatenated traces were bandpass-filtered (80 - 150 Hz, Chebyshev 4th order filter, function filtfilt.m in Matlab) and the voltage values during periods ± 75 msec around the trial onsets/offsets were set to zero, to avoid the edge effects resulting from filtering discontinuous traces. The analytical amplitude was obtained by computing the absolute value of Hilbert-transformed filtered trace (function hilbert.m in Matlab) and z-scored (Extended Data Fig. 2a). Detected events were considered aSWRs if the z-scored analytical amplitude remained above the lower threshold (z = 2) for 20 - 100 msec and if the peak value during this period exceeded higher threshold (z = 5). Only the channels with >150 detected aSWR events were used in the analysis. If the multiple channels from a single subject passed this criteria, a channel with highest number of detected aSWRs was selected for further aSWR-related analysis. Due to the low number of detected aSWRs, one subject was eliminated from the aSWR-related analysis.

### Unsupervised decomposition of iEEG signal

To assess the memory reinstatement, high-frequency activity (HFA; 30-280 Hz) was used as an indirect measure of local populational activity^6–9^. To avoid the effect of low-frequency harmonics on the HFA estimate, we applied the Ensemble Empirical Mode Decomposition^7,10^ (EEMD; https://github.com/leeneil/eemd-matlab.git). Briefly, the EEMD decomposes a non-stationary signal into its elementary components, referred to as intrinsic mode functions^10^ (IMFs; Extended Data Fig. 6). The procedure iteratively applies an empirical mode decomposition algorithm, while adding white noise to prevent the mode mixing^10,11^. Using this approach, decomposition output entirely depends on the signal’s intrinsic properties, avoiding prior assumptions^7,10,11^. The resulting IMFs captured several canonical spectral features consistently across subjects and anatomical structures (Extended Data Table 2). Finally, the HFA time-series on individual channels were reconstructed by summing the channel-specific IMFs with center frequencies > 30 Hz^7^.

### Time-frequency representation of the HFA

The instantaneous spectral power at each time-frequency bin was derived from the reconstructed HFA time series (*x*), using a wavelet transform^12,13^. This approach consists of convolving the time series *x*with a set of Morlet wavelets, parametrized by a range of cycle numbers (*n* = 2, 3, …, 10) at a given frequency *f*,

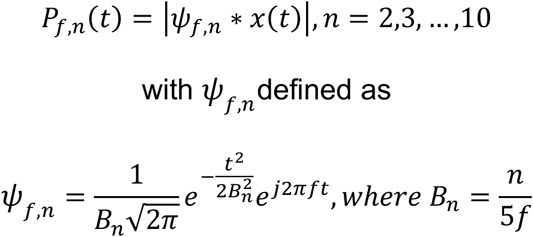

and computing the geometric average 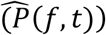 of resulting spectral power at each time-frequency bin:

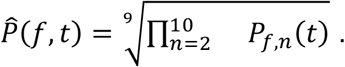

This approach results in a high temporal and frequency resolution, facilitating the detection of narrow-band, transient oscillatory events^12,13^. The wavelet center frequencies were within 30 - 280 Hz range, with 1 Hz increments. The wavelet cycle number range (2-10) is commonly used^14^. To avoid the edge effects, this procedure was applied on the entire individual recording sessions, and the resulting time-frequency response matrices were segmented into trial epochs (starting -1000 msec prior to stimulus onset and ending 1000 msec after the response time). The power within each trial epoch was then normalized by z-transforming each frequency bin and subtracting the average pre-trial baseline (−1000 - 0 msec, relative to stimulus onset^14^).

### Representational Similarity Analysis (RSA)

The representational similarity was quantified as the Spearman correlation between the HFA power spectral vectors (PSVs), for each combination of the encoding-response time bins from the same trial^15–18^ (Extended Data Fig. 6). Specifically, the instantaneous spectral power at each frequency was estimated for 100 msec time bins (10 msec step size, 90% overlap), producing the time bin - specific power spectrum vectors (PSV), spanning the encoding (2 sec time window after stimulus onset) and post-encoding response (time window after stimulus offset and before button press) periods:

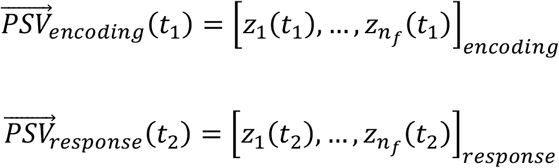

Similar to previous studies^15–20^, we computed Spearman’s correlation as a measure of PSV similarity between the encoding time *t*_1_ and response time *t*_2_ for each encoded stimulus,

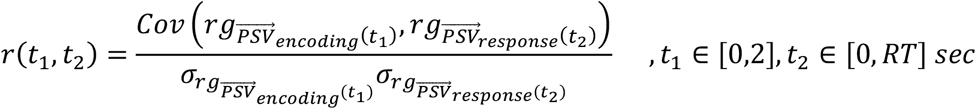

with *rg* representing the ranking operator on the vector 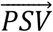, and *σ* the variance of the vector. This produced a trial-specific two-dimensional similarity matrices, containing all the combinations of encoding (*t*_1_) and response (*t*_2_) time bins (Extended Data Figure 6d). The correlation coefficients *r* were then Fisher transformed, with the resulting coefficients following Gaussian distribution. The region-specific (amygdala and hippocampus) similarity matrices were averaged across trials within individual subjects, and used for group-level statistical analysis.

### aSWR-locked memory reinstatement

Memory reinstatement during individual post-encoding time bins was computed by averaging the bin-specific similarity with the encoding period (200 time bins over 2 sec), resulting in a memory reinstatement time series. To obtain the aSWR-locked memory reinstatement, we averaged the memory reinstatement within ± 250 msec around the individual aSWR peak times, separately for amygdala and hippocampus (Fig. 3a). We next tested whether the memory reinstatement is locked to aSWRs (Fig. 3b), by comparing the grand-average aSWR-locked reinstatement trace with an empirical null distribution obtained from Monte Carlo simulation. Specifically, we circularly randomly jittered the aSWR peak times within ± 500 msec window for 1000 times, obtaining an empirical null distribution of memory reinstatement strength.

To test whether the aSWR-locked reinstatement is associated with stimulus-induced arousal and later discrimination (Fig. 3c), we first derived the aSWR-triggered reinstatement, a metric taking the time-locked specificity relative to aSWR peak time into account. For every per-aSWR reinstatement trace around aSWR peak time, we circularly jittered the time as the procedure described above. This results in an empirical null distribution of reinstatement (i.e., correlation coefficient) for every time point around aSWR. We normalized the real reinstatement by z-scoring with mean and standard deviation of the null distribution. We referred to the resulting z-value as aSWR triggered reinstatement and it follows Gaussian distribution. We quantified the aSWR-locked reinstatement difference between the high/low arousal and between correct/incorrect discrimination at every time point by t-test, and corrected for the multiple comparisons using cluster-based nonparametric permutation test. Specifically, we performed the group-level comparisons using paired t-test and identified contiguous time bins with the p < 0.05, defined as clusters. The t-values within each cluster were summed as the cluster statistics. We created an empirical null distribution by shuffling the conditional trial labels 1000 times where the maximum cluster statistics was identified for each permutation. It is considered as statistically significant if the real t-sum cluster statistics exceeded the 95% percentile of the null distribution.

### Cross-structure joint aSWR-locked memory reinstatement

The cross-structure joint aSWR-locked memory reinstatement was obtained by calculating the outer product between the structure-specific reinstatement traces (hippocampus and amygdala) during post-encoding aSWR windows. The resulting joint reinstatement matrices were averaged across the individual aSWRs for each subject, separately for later correctly or incorrectly discriminated trials. To assess the statistical significance of joint cross-structure memory reinstatement, we performed a Monte Carlo simulation to generate an empirical null distribution by circularly jittering the aSWR peak times. The reinstatement significance was defined as exceeding the 95% percentile of null distribution (Fig. 3d).

### Dual states analyses

Recorded periods were divided into low-and high-theta (3 - 10 Hz) or gamma (30 - 250 Hz) periods, based on the subject-specific power median split. The aSWR occurrences are defined as the proportions of aSWRs occurring during each period. The aSWR occurrence comparisons between the low- and high-theta or gamma periods were performed using one-tailed Wilcoxon signed-rank test (p < 0.05; Extended Data Figure 9).

### Mutual information

Mutual information (MI)^14,21^ is a method for quantifying the amount of information shared between the variables of interest. In electrophysiology, MI is applied to test for the presence and directionality of information flow between the multiple time-series. We applied MI to assess the directional influence between the memory reinstatement in amygdala and hippocampus during the post-encoding aSWR windows (Fig. 3e). First, the structure-specific memory reinstatement traces from the amygdala and hippocampus were obtained around each aSWR event (± 250 msec; see aSWR-locked memory reinstatement). Next, we calculated the MI between the amygdala and hippocampal memory reinstatement traces, using the 200 msec bin size (10 msec step size), covering the ± 250 msec window around aSWR peaks. For each time bin, the reinstatement strength was binned into 10 bins (with uniform bin count), consistently across the subjects and conditions. The MI between the time series X and Y was defined as

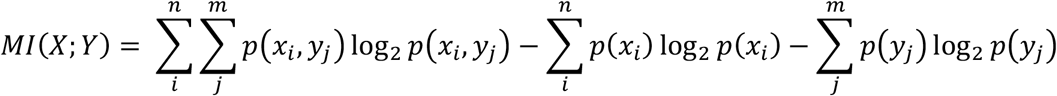

where *p*(*x*_*i*_) and *p*(*y*_*j*_) represented the marginal probability of signals X and Y, *p*(*x*_*i*_, *y*_*j*_) indicated their joint probability, while m and n represented the numbers of reinstatement strength bins for time series X and Y^14,21^. To test the directionality of information flow, we calculated the time-lagged MI by shifting one time series relative to another across all the time bin combinations. The *MI*_*AMY* → *HPC*_ and *MI*_*HPC* → *AMY*_ at individual time bins were defined as the mean of all the subsequent time-lagged MI bins in the other region^14,22^. We defined the MI directional influence as the significant difference between the *MI*_*AMY* → *HPC*_ and *MI*_*HPC* → *AMY*_, assessed using Wilcoxon signed-rank test for each time bin. Correction for multiple comparisons was performed using the cluster-based nonparametric permutation test.

## References

1. Cahill, L., Mcgaugh, J.L. & Cahill, L. 2236, 22983–22986 (1998).

2. Kensinger, E.A. Emot. Rev. 1, 99–113 (2009).

3. Szőllősi, Á. & Racsmány, M. Mem. Cogn. 48, 1032–1045 (2020).

4. Talmi, D. Curr. Dir. Psychol. Sci. 22, 430–436 (2013).

5. Yonelinas, A.P. & Ritchey, M. Trends Cogn. Sci. 19, 259–267 (2015).

6. Ben-Yakov, A., Eshel, N. & Dudai, Y. J. Exp. Psychol. Gen. 142, 1255–1263 (2013).

7. Sols, I., DuBrow, S., Davachi, L. & Fuentemilla, L. Curr. Biol. 27, 3499–3504.e4 (2017).

8. Logothetis, N.K. et al. Nature 491, 547–553 (2012).

9. Skelin, I. et al. Proc. Natl. Acad. Sci. U. S. A. 118, (2021).

10. Buzsáki, G. Hippocampus 25, 1073–1188 (2015).

11. Wu, C.T., Haggerty, D., Kemere, C. & Ji, D. Nat. Neurosci. 20, 571–580 (2017).

12. Jadhav, S.P., Kemere, C., German, P.W. & Frank, L.M. Science (80-.). 336, 1454–1458 (2012).

13. Leal, S.L., Tighe, S.K. & Yassa, M.A. Neurobiol. Learn. Mem. 111, 41–48 (2014).

14. Zheng, J. et al. Neuron 102, 887–898.e5 (2019).

15. McGaugh, J.L. Annu. Rev. Psychol. 66, 1–24 (2015).

16. Bragin, A., Engel, J., Wilson, C.L., Fried, I. & Buzsáki, G. Hippocampus 9, 137–142 (1999).

17. Genzel, L. et al. Philos. Trans. R. Soc. B Biol. Sci. 375, 4–6 (2020).

18. Wixted, J.T. et al. Proc. Natl. Acad. Sci. U. S. A. 111, 9621–9626 (2014).

19. Lopes-dos-Santos, V. et al. Neuron 100, 940–952.e7 (2018).

20. Girardeau, G., Inema, I. & Buzsáki, G. Nat. Neurosci. 20, 1634–1642 (2017).

21. Trouche, S., Pompili, M.N. & Girardeau, G. Curr. Opin. Physiol. 15, 230–237 (2020).

22. Joo, H.R. & Frank, L.M. Nat. Rev. Neurosci. 19, 744–757 (2018).

23. McGaugh, J.L. Proc. Natl. Acad. Sci. U. S. A. 110, 10402–10407 (2013).

## References

1. Maris, E. & Oostenveld, R. J. Neurosci. Methods (2007).doi:10.1016/j.jneumeth.2007.03.024

2. Leal, S.L., Tighe, S.K. & Yassa, M.A. Neurobiol. Learn. Mem. 111, 41–48 (2014).

3. Peirce, J.W. Front. Neuroinform. 2, 1–8 (2009).

4. Avants, B.B. et al. Neuroimage 54, 2033–2044 (2011).

5. Oostenveld, R., Fries, P., Maris, E. & Schoffelen, J.M. Comput. Intell. Neurosci. 2011, (2011).

6. Ray, S. & Maunsell, J.H.R. PLoS Biol. 9, (2011).

7. Lopes-dos-Santos, V. et al. Neuron 100, 940–952.e7 (2018).

8. Wixted, J.T. et al. Proc. Natl. Acad. Sci. U. S. A. 111, 9621–9626 (2014).

9. Canolty, R.T. & Knight, R.T. Trends Cogn. Sci. 14, 506–515 (2010).

10. Wu, Z. & Huang, N.E. Adv. Adapt. Data Anal. 1, 1–41 (2009).

11. Huang, N.E. et al. Proc. R. Soc. A Math. Phys. Eng. Sci. 454, 903–995 (1998).

12. Moca, V. V., Bârzan, H., Nagy-Dăbâcan, A. & Mureşan, R.C. Nat. Commun. 12, 1–18 (2021).

13. Bârzan, H. 2220–2224 (2020).

14. Cohen, M.X. MIT Press (2014).

15. Yaffe, R.B. et al. Proc. Natl. Acad. Sci. U. S. A. 111, 18727–18732 (2014).

16. Lohnas, L.J. et al. Proc. Natl. Acad. Sci. U. S. A. 115, E7418–E7427 (2018).

17. Zhang, H., Fell, J. & Axmacher, N. Nat. Commun. 9, (2018).

18. Norman, Y. et al. Science (80-.). 365, (2019).

19. Pacheco Estefan, D. et al. Nat. Commun. 10, (2019).

20. Staresina, B.P. et al. Elife 5, 1–18 (2016).

21. Quian Quiroga, R. & Panzeri, S. Nat. Rev. Neurosci. 10, 173–185 (2009).

22. Helfrich, R.F. et al. Nat. Commun. 10, 1–16 (2019).

